# Green Leaves, Red Flags: Host Plant Associations as Indicators of Prey Defense for an Avian Predator

**DOI:** 10.64898/2026.09.27.754804

**Authors:** Nich W Martin, Kathryn E Sieving, Bryan M Kluever, Jaret C Daniels, Heather J McAuslane

**Affiliations:** Department of Entomology and Nematology, University of Florida, Gainesville, FL, USA; Florida Museum of Natural History, University of Florida, Gainesville, FL, USA; Department of Wildlife Ecology and Conservation, University of Florida, Gainesville, FL, USA; US Dept of Agriculture, Animal and Plant Health Inspection Service, Wildlife Services, National Wildlife Research Center, Gainesville, FL, USA; School of Natural Resources and Environment, Institute of Food and Agricultural Sciences, College of Agricultural and Life Sciences, University of Florida, Gainesville, FL, USA; McGuire Center for Lepidoptera and Biodiversity, Florida Museum of Natural History, University of Florida, Gainesville, FL, USA

## Abstract

Defended prey are often thought to rely on conspicuous warning signals to distinguish themselves from undefended prey, facilitating predator learning and avoidance. Yet many chemically defended organisms are inconspicuous and difficult to distinguish from palatable prey, raising a fundamental question: how do predators avoid defended prey when prey themselves provide little or no reliable signals of defense? We tested whether host plants can function as indirect indicators of prey defense using Carolina wrens (*Thryothorus ludovicianus*), chemically defended *Eumaeus atala* larvae, undefended *Galleria mellonella* larvae, and two distinct plant species. Defended and undefended prey were experimentally made to appear visually identical, forcing predators to rely on host-plant associations as the only available source of information. Wrens rapidly altered both search behavior and attack decisions according to plant identity. Mechanistic attack models incorporating predator discrimination between defended and undefended prey were strongly favored when plants were present, with defended prey experiencing substantially lower attack risk than undefended prey. When plants were removed, support for discrimination weakened and attacks on both prey types increased. Spatial autocorrelation of host plants further shaped predator behavior, with strongly clustered plants hosting defended prey receiving disproportionately fewer visits. Together, these results suggest host plants can function as ecologically meaningful cues of prey defense, influencing both predator search patterns and attack decisions. More broadly, our findings suggest warning systems may extend beyond prey phenotype alone, and that ecological context—including host-plant identity and spatial structure—may contribute to the evolution and maintenance of aposematism.

## Introduction

Natural enemies encounter a myriad of different prey types, many of which are defended in one way or another. Because of the selective pressure prey defense places on natural enemies, experienced predators make strategic decisions about what prey to consume (Barnett et al., 2012). In turn, highly defended prey tend to have conspicuous signals (Poulton, 1890), a phenomenon called aposematism, which is thought to better facilitate predator learning by capitalizing on their strategic decisions, providing a robust signal that allows defended prey to avoid predation (Guilford, 1990). Predator learning is therefore central to the evolution and effectiveness of aposematism. Theoretical models have long predicted that the evolution of warning coloration depends not only on the costs and benefits of defense itself, but also on how predators learn to associate prey characteristics with unprofitability (Leimar et al., 1986). Importantly, conspicuousness may facilitate avoidance learning, but warning traits can also be favored because they allow predators to reliably discriminate defended from undefended prey (Sherratt, 2002; Sherratt and Beatty, 2003). Thus, the effectiveness of a warning system depends on the information available to predators and their ability to use that information when making foraging decisions.

Despite extensive research on visually conspicuous warning signals and predator learning, comparatively less attention has been given to defended prey that remain visually inconspicuous and difficult to distinguish from palatable individuals (however, see Skelhorn and Rowe, 2010). This is despite the existence of visually inconspicuous, defended prey (Paul and Van Alstyne, 1988; Santos et al., 2003; Dossey et al., 2012) and theory predicting that as populations evolve from inconspicuous, undefended phenotypes to aposematic forms, transitional phenotypes of inconspicuous, defended prey should exist (Ruxton et al., 2019). Understanding these dynamics is non-trivial, as mixed populations of defended and undefended individuals with no distinguishing signals present a problem for predators. Specifically, when signals distinguishing defended and undefended individuals are absent, predator fitness is predicted to decrease (Sherratt, 2003; Holen et al, 2012).

The absence of a distinctive visual phenotype, however, does not necessarily mean that predators lack information about prey profitability. Predators can learn to use environmental or contextual cues that predict the profitability of prey, allowing the same prey phenotype to elicit different responses in different contexts. For example, great tits (*Parus major*) learned to associate prey palatability with environmental context and subsequently altered their preferences for the same prey type depending on the context in which it was encountered (Hansen et al., 2010). Such contextual learning provides a potential mechanism by which predators could discriminate among otherwise similar prey when information about their profitability is associated with features of the environment rather than with the prey phenotype itself. Moreover, if prey defenses were closely associated with environmental stimuli that were reliably consistent through time and space, then sufficiently sophisticated predators could learn to strongly avoid defended prey which are otherwise identical to undefended individuals. Such associations could influence not only whether predators attack a prey item after encountering it, but also where predators search for prey in the first place. Optimal foraging theory predicts that predators should balance the energetic costs of searching and moving among potential feeding sites against the expected profitability of those sites (Stephens and Krebs, 1986). When environmental features reliably predict prey profitability, predators should therefore benefit from incorporating those features into decisions about where to allocate search effort. The spatial distribution of such environmental cues should consequently matter: when cues associated with defended prey are spatially clustered, predators may be able to avoid relatively large areas after learning their association, whereas weakly structured environments may require more frequent discrimination among potential prey encounters.

The scenario described above is highly relevant to one particular group of organisms, soft-bodied insects who obtain their defenses by sequestering and storing chemical toxins synthesized by their host plants. In many cases, these insects are highly specialized, feeding on a small set of closely related, morphologically similar plant species. Recent work suggests antipredator defenses should not necessarily be understood through a single sensory modality or as isolated traits expressed by the prey itself. Instead, prey defenses can operate across multiple stages of the predation sequence and can involve combinations of visual, chemical, behavioral, and environmental information (Kikuchi et al., 2023). Host plants may therefore provide predators with information about prey profitability.

Here we tested the hypothesis that plants could serve as prey defense signals through a set of controlled experiments using a small, insectivorous avian predator, larvae of two lepidopteran species, one chemically defended and the other undefended, and two morphologically distinct plant types. We did this by masking defended and undefended prey to make them appear identical to each other, making host plant characteristics the only signals available to predators. If this hypothesis is valid, predators should learn to associate specific plant types with defended or undefended prey. We tested predator ability to make this association by observing both the probability that a predator returned to plants hosting defended prey as well as the attack rates on defended and undefended prey when plants were present vs. absent.

## Methods and Materials

### Prey

We used *Eumaeus atala* Poey (Lycaenidae) third to fourth instar larvae as the defended prey, which gains its defenses through sequestration of azoxyglycoside compounds from its host plant, *Zamia integrifolia* Linnaeus f. (Zamiaceae) (Rothschild et al., 1986). These compounds are well documented to have negative effects on both predator behavior (Bowers and Larin, 1989; Bowers and Farley, 1990) and physiology (Healy, 1969). Larvae were reared from eggs laid by adults, which were collected from the Montgomery Botanical Center, Coral Gables, Miami-Dade County, Florida. Larvae were provided fresh *Z. integrifolia* leaflet cuttings *ad libitum* until they reached their third or fourth instar, such that they were approximately the same size as specimens of the undefended prey species.

For the undefended prey, we used greater wax moth third instar larvae (*Galleria mellonella* Fabricius [Pyralidae]) which are known to be highly palatable to many insectivores (Strzelewicz et al., 1985). Wax moth larvae were purchased from an exotic pet store in Gainesville, Florida, and were kept in a refrigerator at 3.75 degrees Celsius until experiments began.

### Plants

We used twenty 6-year-old *Z. integrifolia* plants in 1-gallon pots, purchased from a local nursery in Gainesville, FL, and branch clippings of laurel oak trees (*Quercus laurifolia* Michx) placed in soil-filled 1-gallon pots. *Zamia* plants were watered once a week, as well as the day before experimentation. One-gallon pots with laurel oak clippings were also watered the day before experimentation.

### Predators

We selected Carolina wrens (*Thryothorus ludovicianus* Latham) as the predator owing to their high degree of insectivory (Gill and Haggerty 2012) via foliage and surface gleaning and lack of neophobia in foraging contexts (Stanbeck and Burke 2020). A total of 42 Carolina wrens (*Thryothorus ludovicianus* Latham) were captured via mist netting and housed individually in large, 6-meter-tall outdoor aviary cages (area: 3.66 x 4.57 m) at the US Department of Agriculture (USDA) Animal Plant Health Inspection (APHIS) Wildlife Services (WS) National Wildlife Research Center (NWRC) Florida Field Station in Gainesville, Florida, USA. Each cage contained a small plot of high shrub vegetation and was equipped with a feeding station and timed sprinkler system. In addition to several nest boxes, vegetation served as refuges for the wrens during, and between, experimental phases. Once captured, wrens were allowed to acclimate for 2 days during which they were given water and dry food (Nesting Superblend®) *ad libitum*, both of which were placed on the feeding station. Wrens were also provided a small tray of live mealworms daily. In addition to the tray, two live mealworms were individually encased in a masking mold which consisted of one part beef tallow (EPIC Provisions), one part water, two parts white flour. From this blend, 237 ml were mixed with approximately 7.5 ml of green food coloring. These molds were later used to conceal differences between defended and undefended prey and were used during this acclimation period to train wrens to associate these molds with live prey (Figure 1). Birds were released back into the wild after completing all experimental trials.

**Figure 1.**
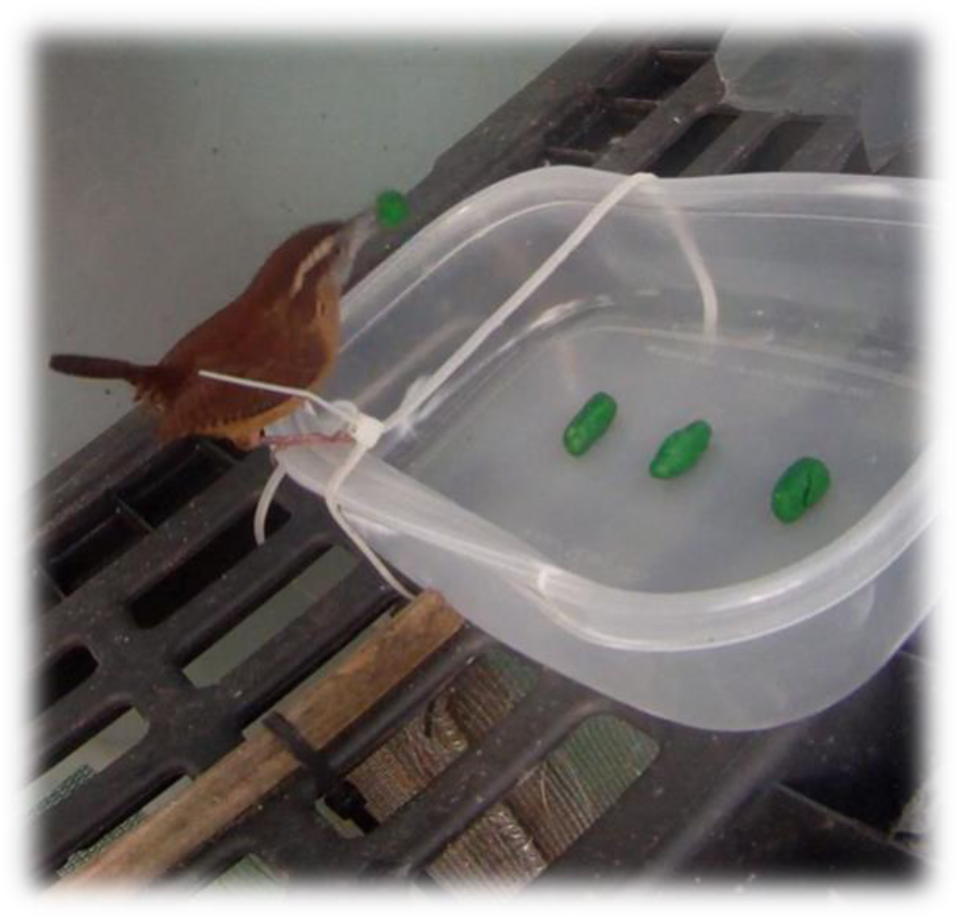
Showing prey in pastry dough molds during the plants absent experimental trial.

### Experimental Design

Individual wrens went through four different phases (Figure 2a). The first was the aforementioned 2-day acclimation phase, where wrens were allowed to adjust to their surroundings and prepare for the experiment. The second was the training phase in which wrens were to make their initial associations between different plant types and the prey types those plants hosted. The third and fourth phases consisted of an experienced (post-training) phase with plants, followed by an experienced phase without plants. We did not randomize the order of these last two phases out of concern that if wrens were presented prey without signals (i.e., without plants) it would disrupt the associations made during the training phase due to so-called “recency effects” (Henne et al., 2021). This, among other concerns, could have caused wrens to terminate foraging when plants were present during the subsequent phase.

**Figure 2a.**
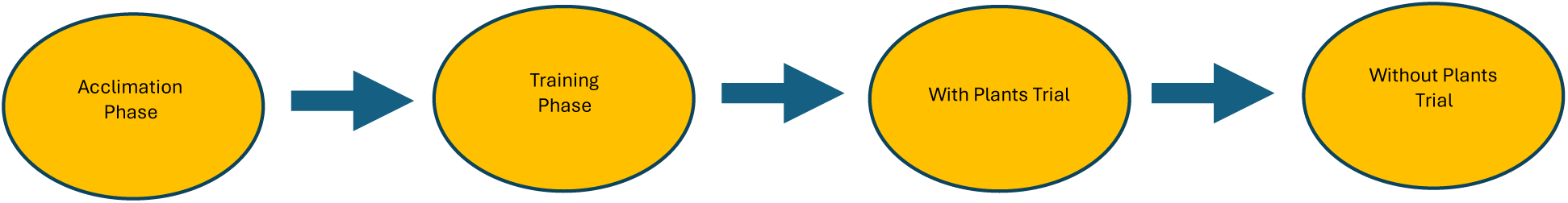
Schematic diagram of different experimental phases.

**Figure 2b.**
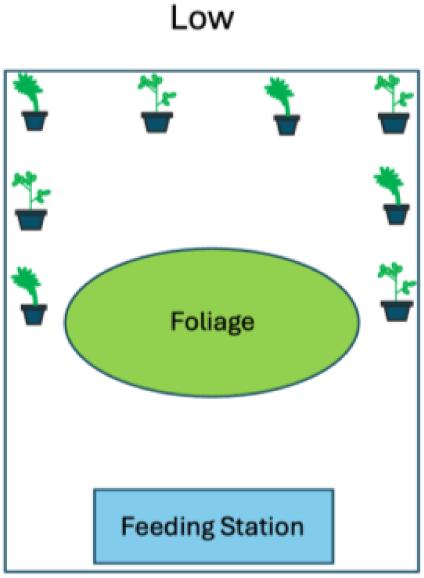
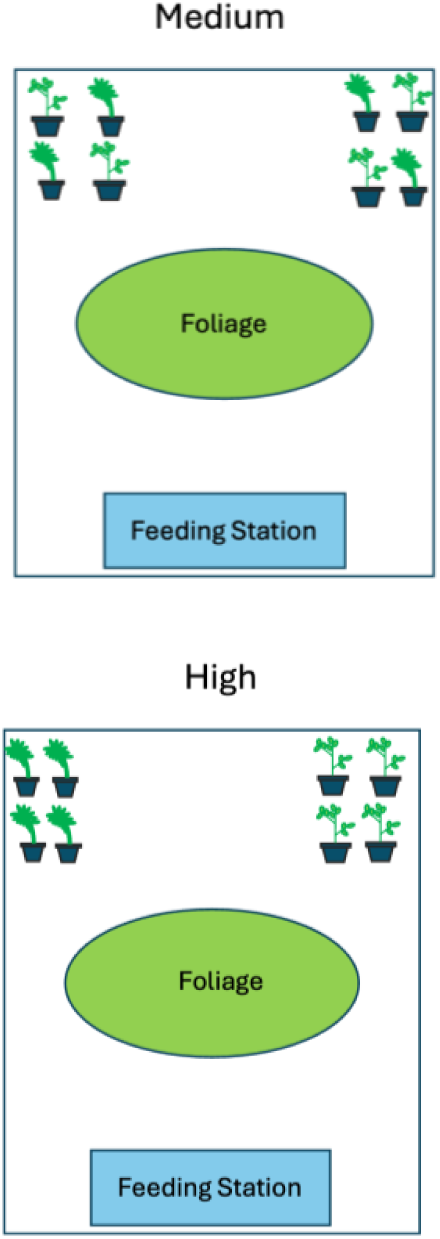
Aviary setup for three different levels (Low, Medium, and High) of plant spatial autocorrelation for two different plant species; single leaflet plants representing *Zamia* plants and three leaflets representing *Quercus*.

We assumed energetic costs exist for moving from one plant to another. Optimal foraging theory would predict predators to minimize their movements to plants known to host low-quality prey and maximize movement efforts to plants hosting high-quality prey. These effects should increase as the energetic cost of moving to (or avoiding) a particular plant type decreases, such as when plants show high spatial autocorrelation. Additionally, some plants which host defense-sequestering herbivores are highly modular (Malcolm et al., 1989), showing high degrees of spatial autocorrelation. As such, large scale size differences between plant species may be just as relevant as finer scale differences. Therefore, during both the training phase and the experienced phase (plants present), plants were randomly assigned one of three different levels of spatial autocorrelation (SAC): “low”, “medium”, and “high”. Plants assigned “high” spatial autocorrelation were clustered together in one corner of the aviary. Plants hosting defended larvae with “medium” spatial autocorrelation were grouped together in pairs of two alongside two plants hosting undefended larvae. Plants with “low” spatial autocorrelation, were spread evenly apart, alternating between plants hosting defended prey and plants hosting undefended prey (Figure 2b).

During the acclimation phase, four plants of each species were placed in the aviary according to their assigned spatial autocorrelation on the side opposite the feeding station. On the day of the training phase, all food was removed from the feeding station 2 hours prior to data collection. Two Nabi® SD19 cameras (Model no. CAMERA2-06- SU13) were strategically placed on small wooden stands in the aviary to simultaneously monitor all plants. Both species of live prey were encased in the same masking mold compound used in the acclimation phase and placed on a randomly assigned plant type. Four to six prey (either defended or undefended) were placed on a single plant of the assigned type. The number of prey varied between replicates due to availability, however, there was an equal number of each prey species for a given replicate. For the training phase, we chose to place all prey of each type on a single plant of the assigned species to account for spatial learning on the part of each predator. We reasoned this would prevent wrens from making spatial associations going into the next phase, with wrens relying solely on plant features.

Cameras were started and the data collection period for the training phase began once the experimenter left the aviary. We observed wrens attacking prey *ad libitum* for 30 minutes. Prey was said to be attacked if a wren either removed a masking mold from the plant or pulled prey out of the mold. After the first attack of a specific prey species, we recorded the subsequent number of times wrens returned to the same species of plant, and the subsequent number of attacks on that prey species. Wrens were said to have returned to a plant if they landed on a leaf, branch, soil inside the garden pot, or the rim of the garden pot. Only wrens which attacked both prey species continued to the next phase (training phase with plants; 28 wrens total). At the end of this training phase, dry food was returned to the feeding station, without live mealworms.

The next day, all food was removed from the aviary 2 hours prior to the start of the next phase. The same number of masked prey from the previous trial was assigned to the same plant species from the training phase, which maintained their spatial autocorrelation from the previous phase; however, prey were distributed across all plants of that plant species. Wrens were again allowed to forage *ad libitum* for 30 minutes, and we recorded the number of times a wren visited each plant and the number of attacks on each prey species.

Immediately after this phase of the experiment, all plants and prey were removed from the aviary, and four new masked prey, two defended and two undefended, were placed in a container fixed to the aviary’s feeding station. This was done to observe wren behavior when the potential signal from plants was removed. Wrens were again allowed to forage for 30 minutes. In this final and fourth phase of the experiment, we only recorded the number of attacks on each prey species.

### Statistical Models and Data Analysis

#### Plant Associations and Predator Foraging

To analyze plant visitation during the experimental (with plant) phase, we treated wren behavior as a two-stage process. First, we modeled whether an individual visited any plants during the trial. Second, conditional on at least one visit, we modeled how plant visits were allocated between prey-hosting category (i.e., defended vs. undefended). Because all plants in a single trial belonged to one of two categories, the allocation response was expressed as the number of visits to defended-prey-hosting plants out of the total number of plant visits. Thus, our estimand was the conditional probability that any given plant visit was directed towards a plant species associated with hosting defended prey. We restricted this analysis to the plants-present experimental phase because the training and with-plant-experimental phase differed slightly in structure and measurement. For each wren *i* we calculated the total number of plant visits during the with-plant-experimental trial as:

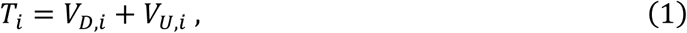

where *V_D_*_,*i*_ is the number of visits to plants of the species hosting defended prey and *V_U_*_,*i*_ is the number of visits to plants of the species hosting undefended prey. We analyzed visitation with a joint Bayesian hurdle model. The first component modeled whether any plant visit occurred,

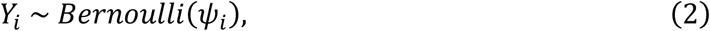

where

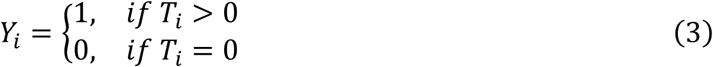

and *ψ_i_* is the probability that wren *i* visited at least one plant. We modeled this on the logit scale, such that,

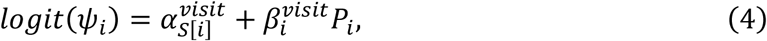

where *α^visit^*is the SAC-specific intercept for the spatial autocorrelation treatment assigned to wren *i* where *S*[*i*] ∈ {*Low*, *Med*, *Hig*ℎ} , *β^visit^* is the effect of experience during the training phase, and *P_i_* is the proportion of prey consumed during the training phase that were defended. The second component modeled allocation of visits provided visiting occurred such that, for all wrens with *T_i_* > 0,

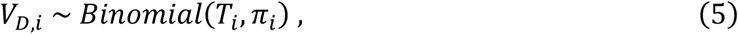

where *π_i_* is the conditional probability that any given visit to a plant was to the species hosting defended prey. We modeled *π_i_* as

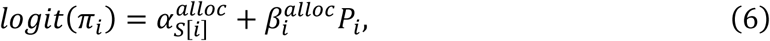

where *α^alloc^* is again the SAC-specific intercept, but this time toward the visit allocation, and *β^alloc^*represents the effect of prior training experience.

For both components, for each bird, the prior experience term *P_i_* was calculated from the training phase as

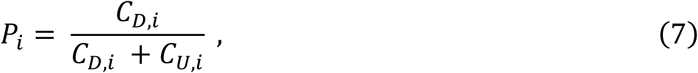

where *C_D_*_,*i*_ and *C_U_*_,*i*_ are the number of defended and undefended prey, respectively, attacked during the training phase. We treated spatial autocorrelation as categorical as opposed to ordinal because we did not want to assume behaviors operated monotonically with degrees of plant clustering, as intermediate levels of spatial autocorrelation could, for example, lead to greater discrimination between plant types.

We assigned weakly regularizing priors to all parameters, with all priors centered on the logit-scale such that,

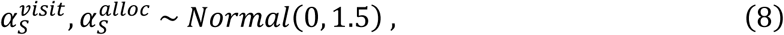

and

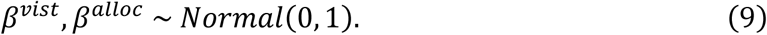

#### Plant Associations and Predator Attacks

For predator attacks, we took a comparative modeling approach, fitting two alternative mechanistic process models to the observed data. We did this for two reasons. First, the temporal execution of our experiments, with non-randomized treatment vs. control trials, lacks statistical independence, making traditional null-hypothesis models untenable. Second, and more importantly, taking an explicit, generative, process modeling approach reveals greater insight into ecological mechanism, allowing for greater scientific inference (McElreath, 2018).

The first model represents attacks when signals are present, where predators can discriminate between defended and undefended prey through some perceptible cue. The second model represents attacks when signals are absent, where predators are incapable of distinguishing prey types. These models provide distinct biological hypotheses about the information available to predators when making attack decisions.

Because both prey types were equally available within trials, we modeled number of attacks on both prey types *A_j_* as binomial outcomes, conditional on the number of prey available for each type,

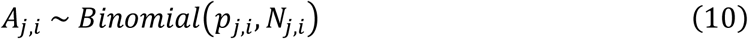

where *A_j_*_,*i*_ is the number of prey type *j* attacked, where *j* ∈ {*defended*, *undefended*}, *N_j_*_,*i*_ is the total number of prey type *j* available, and *p_j_*_,*i*_ represents the realized probability of *j* type prey being attacked. Attacks on defended and undefended prey were analyzed jointly because, under both candidate models, their realized attack probabilities are deterministic functions of the same underlying parameters. Thus, observations from both prey types contribute to inference on a shared mechanistic process rather than being treated as independent, or independence-violating, models.

For the signals absent model, predators cannot tell the difference between prey types. Thus, both defended and undefended prey share the same realized probability of attack. We assumed this probability is determined by the baseline attack rate *α* and the protective effect of defense *θ*. This effect is attenuated by the relative abundance of undefended prey, which is due to the positive reinforcement predators experience attacking and consuming undefended prey. Thus, the protective effect of defense weakens as the perceived proportion of defended prey, *P*, decreases. Like Broom et al., (2005), we assumed the effect of defense *θ* on baseline attack rates decays exponentially, such that

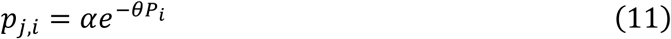

where *P_i_* is the proportion of the prey population wren *i* perceives to be defended, and was calculated according to equation 7.

The signals present model has a similar structure for defended prey but allows predators to discriminate between prey types. Therefore, the probability of attack on defended prey remains

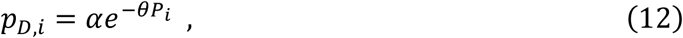

while the probability of attack on undefended prey is simply a function of baseline attack rate,

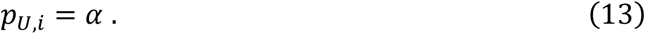

Because *p_D_* and *p_U_* differ in structure under our signals present model, the realized probability of being attacked is conditional on prey type such that,

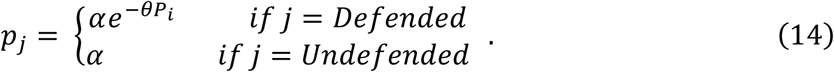

Because attack probabilities represent the likelihood of an individual being attacked, we restricted *α* to the interval (0,1) and assigned it a weakly regularizing normal prior transformed by the inverse logit function, such that,

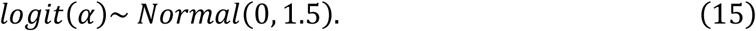

We assumed defenses necessarily reduce attack probability, not increase it. Therefore, we assigned *θ* a half-normal prior with a mean of 0 and a standard deviation of 1.

We used predictive simulation from the joint posterior distributions to estimate *p_j_* and generate posterior contrasts for the best-supported models under plant-present and plant-absent conditions. We compared the relative support for the different models using log-transformed Bayes factor, calculated as the log ratio of marginal likelihoods, which were estimated by bridge sampling via the bridgesampling package (version 1.1-2) in R. We conducted model comparisons separately for datasets collected when plants were present and when plants were absent. Hamiltonian Monte Carlo was performed with Stan (version 2.37) using the rstan package (2.36.0.9000). All posterior simulations and figures were generated either using base R (4.2.3) or the ggplot2 package (4.0.1) with an R studio (2023.09.1) interface.

## Results

### Plants Visited

Most wrens visited at least one plant during the experimental phase, with spatial autocorrelation having no meaningful effect on whether wrens visited any plants. In contrast, spatial autocorrelation had a noticeable effect on whether plant visits were directed to a species hosting defended prey. Posterior mean probabilities of visiting a plant hosting defended prey were lowest under high SAC (with 0.41 90% CI: 0.31 to 0.51 for low, 0.50 CI: 0.43 to 0.57 for medium, and 0.17 CI: 0.12 to 0.25 for high SAC).

The proportion of defended prey attacked during the training phase (training-effect), had little influence on whether wrens visited any plants in the experimental phase (41% chance of a negative slope), suggesting toxin load from consuming defended prey was not limiting wrens’ ability or willingness to forage. In contrast, the allocation of visits to plants hosting defended prey showed a predominately (81% chance) negative slope (posterior mean = −0.64, 90% CI = −1.81 to 0.56). While uncertainty was substantial, this corresponds to an estimated decline of approximately 0.08-0.15 in the probability of visits to plants hosting defended prey across the full range of defended prey attacked during the training phase.

### Prey Attacked

#### Plants present

During the experimental phase, when plants were present, the data were in very strong favor of the signals-present model over the signals-absent model (log Bayes factor = 14.55) with a significantly smaller proportion of the available defended prey being attacked compared to when plants were absent (Figure 3A and 3B). Moreover, posterior distributions for the baseline attack parameter *α* (Figure 4A and 4B) and defense-effect parameter *θ* (Figure 4C and 4D) differed strongly between plant-present and plant-absent conditions. Mean baseline attack probability was *α* = 0.52 (90% CI: 0.45-0.59) and the posterior mean for the effect of defense was *θ* = 2.24 (90% CI: 1.57-2.94). Structurally, the realized probability of attack on undefended prey is *α* and therefore equal to 0.52. Sampling across all values of *P* during the training phase, the corresponding probability of attack on defended prey was 0.17 (90% CI: 0.12 to 0.23; Figure 5A). Thus, defended prey are predicted to experience significantly fewer attacks than undefended prey when plants are present (mean difference = −0.34, 90% CI: −0.42 to −0.25).

**Figure 3.**
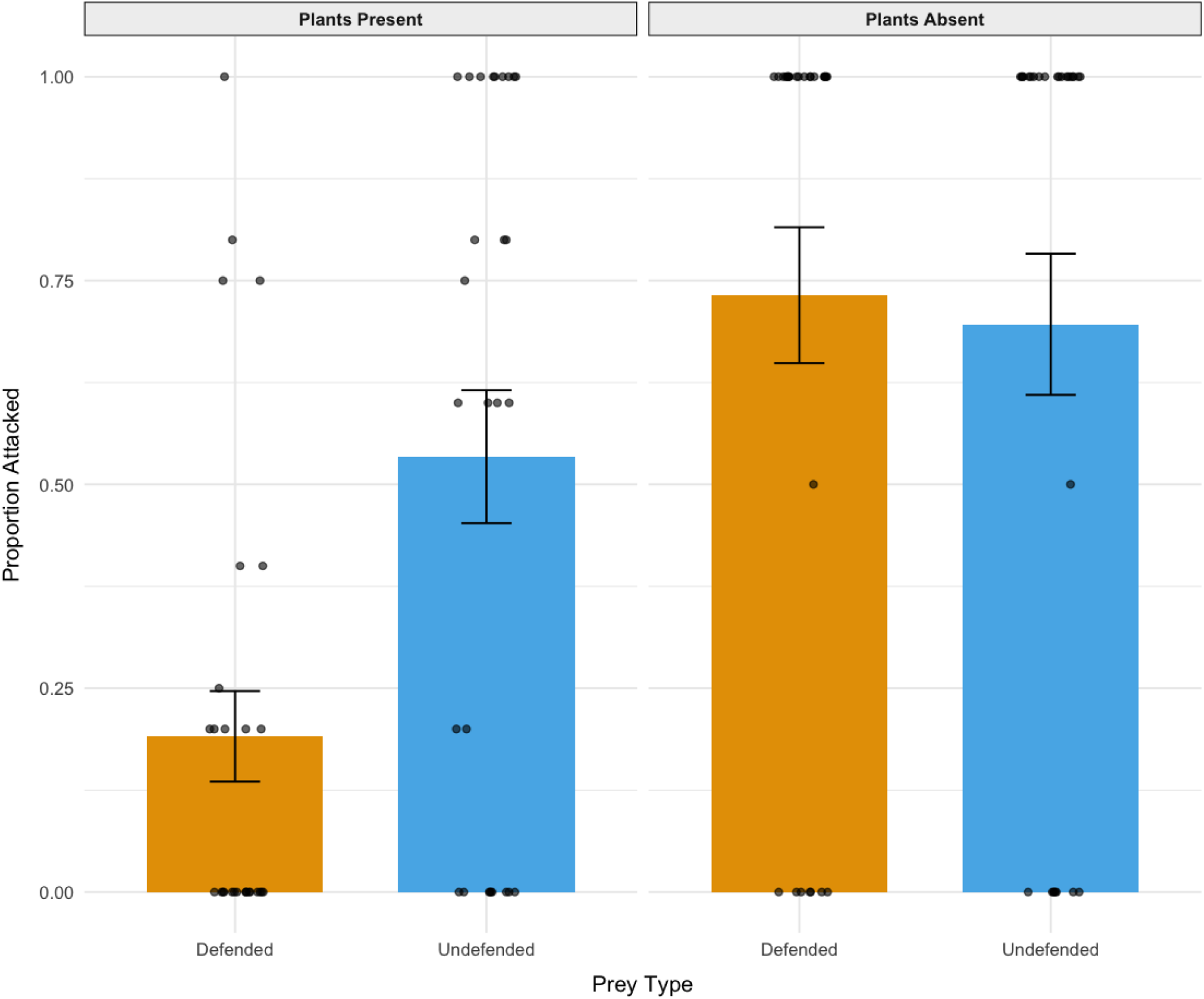
Observed proportions of defended prey (orange) and undefended prey (blue) attacked during the experienced phases with plants and without plants.

**Figure 4.**
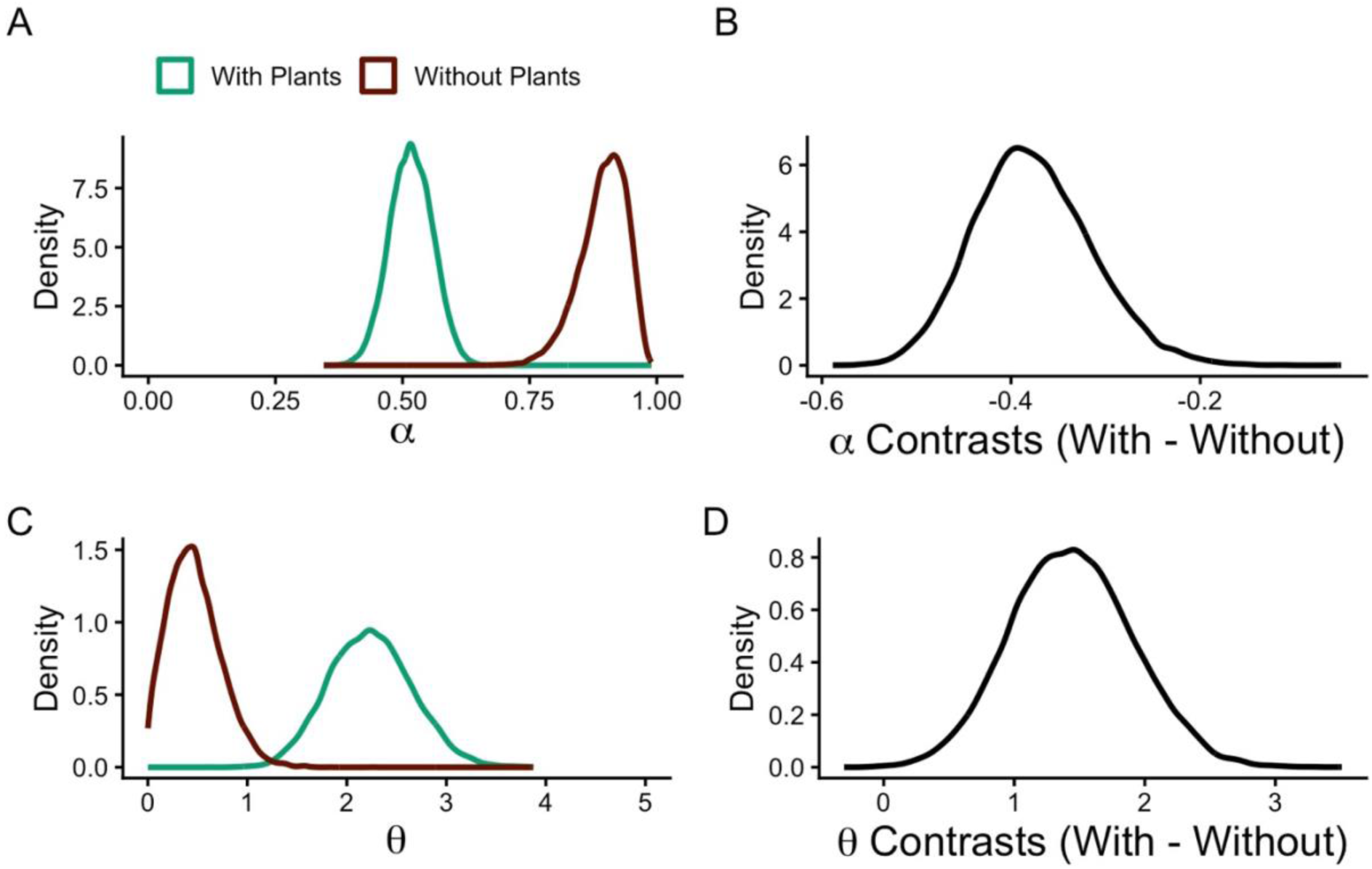
Posterior distribution and contrast density plots for the intrinsic likelihood of being attacked *α* and the inhibitory effect of defense *θ* when plants are present (green) and when plants are absent (red).

**Figure 5.**
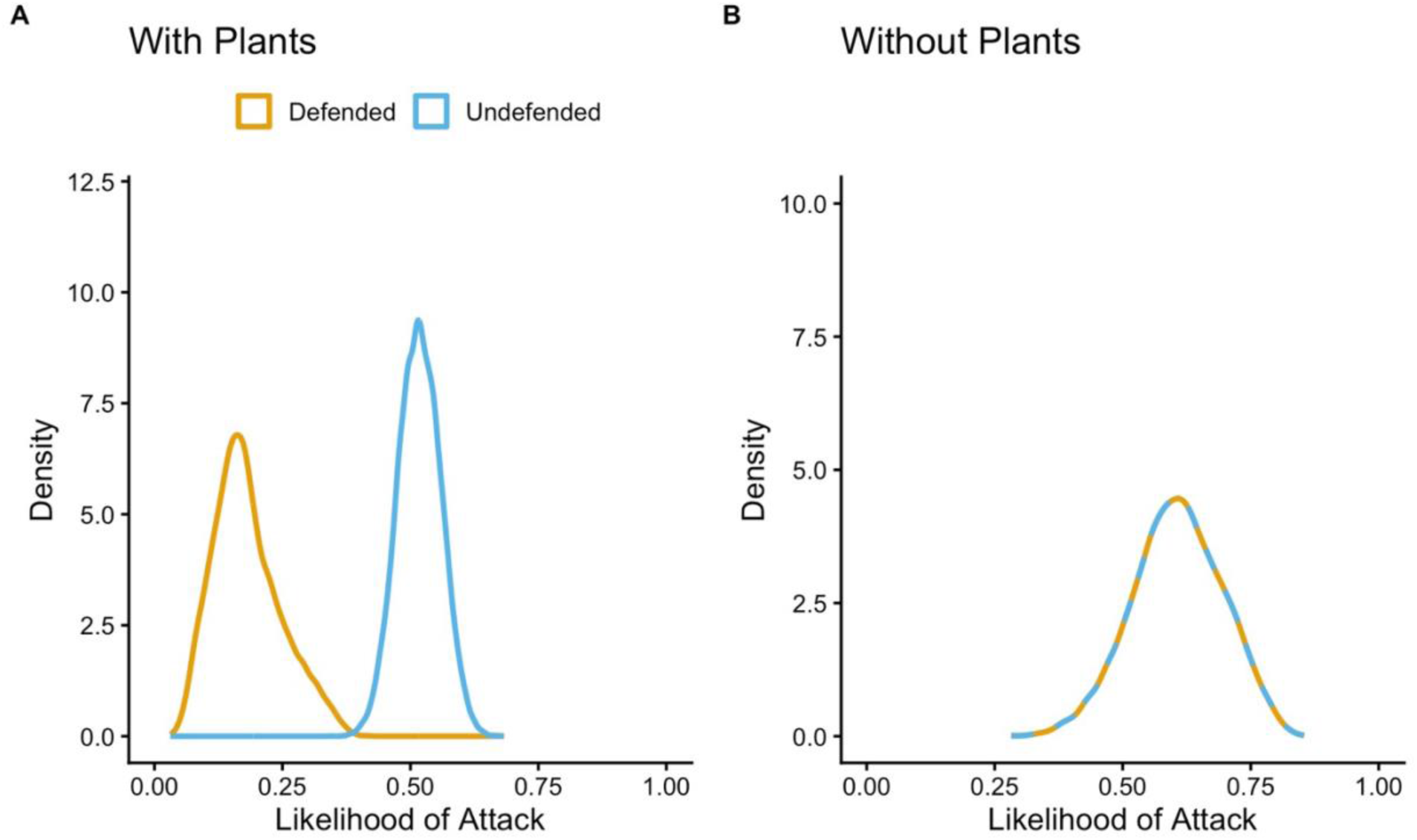
Posterior distribution density of the realized likelihood of being attacked for defended prey (orange) vs. undefended prey (blue) when plants are present vs. absent.

#### Plants absent

When plants were absent, the data favored the signals-absent model (log Bayes factor = −5.49), suggesting strong evidence that wrens were unable to distinguish defended from undefended prey. Additionally, the posterior mean for baseline attack probability was substantially higher than its predicted value when plants were present, mean *α* = 0.89 (90% CI: 0.81 to 0.96). Likewise, defense effect *θ* was much smaller, mean *θ* = 0.8 (90% CI: 0.46-1.15). As such, sampled across *P*, for both the training and experimental phase when plants were present, the realized posterior predicted probability of attack on both prey types was 0.61 (90% CI: 0.46 to 0.75; Figure 5B). Consequently, even with the deterrent effects of *θ* overall attack risk was higher for both prey types when plants were absent.

## Discussion

### Predators associated host plants with defended prey

When defended and undefended prey are difficult to distinguish, undefended individuals are likely protected, while defended individuals experience increased rates of predation (Turner et al., 1984). By linking prey defense to a stable, predictable environmental feature, plant associations may allow predators to avoid defended individuals, absent direct signaling from prey. Our results strongly support this hypothesis, suggesting host plants can function as environmental cues associated with prey defense. When plants were present, attack data very strongly favored the model in which wrens could discriminate between defended and undefended prey, with defended prey experiencing substantially lower realized probabilities of being attacked. When plants were absent, support was in favor of the model in which wrens could not discriminate between prey types, but more weakly so. Moreover, when plants were absent, overall attack rates on both prey types increased, consistent with others’ findings (Jones et al., 2012). More broadly, these results are consistent with evidence that predators can use environmental context to modify their responses to prey (Hansen et al., 2010), and suggest that information about prey profitability need not be encoded solely in prey phenotype. Our visitation results extend this concept beyond individual features of the host plant to include spatial structure of the environment.

Spatial autocorrelation had little effect on whether wrens visited plants at all, but we observed strong effects on the allocation of those visits between plant categories (e.g., defended-prey-hosting vs. undefended-prey-hosting). Specifically, wrens under high SAC treatments (Figure 2) were much less likely to direct visits toward plants hosting defended prey compared to wrens under low and medium SAC treatments. This suggests strong clustering of host plants can alter how predators use plant-associated cues. More broadly, this pattern indicates that plant-associated cues influence predation at more than one stage of interaction. Host plants did not simply affect whether defended prey were attacked once encountered; their spatial arrangement also altered how predators distributed search effort among potential encounter sites. As such, plant features may influence both exposure to predators and the consequence of prey being encountered. This pattern is consistent with the prediction that predators should incorporate the spatial structure of environmental cues into their search decisions when those cues predict prey profitability (Stephens and Krebs, 1986).

Taken together, our attack and visitation analyses suggest host plants may operate as part of an extended warning environment. That is, host plants do not merely co-occur with defended prey, but may also help structure the conditions under which defense becomes ecologically effective. This has potentially significant evolutionary consequences for specialist herbivores, particularly those that derive defenses from their host plants. Rather than suggesting that host plants themselves function as warning signals in the strict sense, our results suggest that the information used by predators to avoid defended prey can be distributed across the ecological context in which prey occur. In this sense, an extended warning environment may complement, instead of replace, prey-borne warning traits.

### Implications for missing steps in the evolution of aposematism

This possibility has interesting implications for the evolution of aposematism. The origin of conspicuous warning signals is often treated as problematic because rare, novel warning phenotypes may initially suffer strong predation before predators learn to avoid them (Speed and Ruxton, 2005). One possible solution would be if some other stable cue in the prey environment already allows predators to associate defense with defended prey. Our results raise the possibility that host-plant associations could serve that function.

If predators can use plant identity as a reliable signal of defense, then prey may receive some protection even before prey-produced warning signals develop. This is likely, as vertebrates readily learn to avoid microhabitats with high predation risk (Wasserlauf et al., 2023; Mathis and Unger, 2012). Likewise, recent works reveal animal avoidance of disease-infected microhabitats, where disgust is a motivating factor (Sarabian et al., 2023). Consuming chemically defended prey is associated with disgust-like reactions (Rozin et al., 2008). These examples further suggest that predators can use environmental characteristics to avoid locations or resources associated with negative consequences, providing a plausible behavioral mechanism by which host-plant associations could influence predation risk. However, whether such associations would facilitate or inhibit the evolution of conspicuous warning traits remains an open question. It should, however, be noted that environmental cues could, in principle, alter selection on warning coloration in more than one direction. If an established host-plant association provides protection to defended prey, it could reduce the initial predation cost of evolving a novel warning phenotype by providing predators with an alternative source of information about defense. Conversely, if predators already reliably avoid defended prey based on host-plant identity, selection for an additional prey-borne warning signal could be reduced. The evolutionary consequence will therefore depend on how reliable the environmental cue is, how strongly predators use it, and how the benefits of an additional warning trait compare with its costs.

Although our experiments were not designed to test the evolutionary origin of aposematism directly, they suggest the effectiveness of prey defense can depend strongly on ecological context. In this study, host plants increased the extent to which defended prey were avoided, and strong spatial clustering altered how predators allocated search effort among plant types. These findings support the view that plant associations can act as meaningful environmental cues of prey defense and suggest that ecological context may play a more active role in the evolution of aposematism than is typically assumed. Future work should focus on whether these learned associations are maintained with the addition of novel, conspicuous signaling.

## Acknowledgments and Funding

Research was conducted under USGS Federal Bird Banding Permit No. 22541, Florida Fish and Wildlife Conservation Commission Permit No. LSSC-20-00022, University of Florida IACUC Protocol No. 201910895, and USDA APHIS Protocol No. QA- 3188.

This research was partially supported by the USDA APHIS Wildlife Services National Wildlife Research Center.

We thank Juliette Rubin and Danyelle Sherman for their invaluable care of Carolina wrens used in this study and Collette St. Mary for her critical feedback on an earlier version of this manuscript.

The findings and conclusions in this publication have not been formally disseminated by the U.S. Department of Agriculture and should not be construed to represent any agency determination or policy.

## Literature Cited

Agrawal, A.A., 2005. Natural selection on common milkweed (*Asclepias syriaca*) by a community of specialized insect herbivores. Evolutionary Ecology Research, 7: 651–667.

Barnett, C.A., Skelhorn, J., Bateson, M. and Rowe, C., 2012. Educated predators make strategic decisions to eat defended prey according to their toxin content. Behavioral Ecology, 23(2), pp.418–424.

Bowers, M.D. and Farley, S., 1990. The behaviour of grey jays, *Perisoreus canadensis*, towards palatable and unpalatable Lepidoptera. Animal Behaviour, 39(4), pp.699–705.

Bowers, M.D. and Larin, Z., 1989. Acquired chemical defense in the lycaenid butterfly, *Eumaeus atala*. Journal of Chemical Ecology, 15, pp.1133–1146.

Broom, M., Speed, M.P. and Ruxton, G.D., 2005. Evolutionarily stable investment in secondary defences. Functional Ecology, 19(5), pp.836–843.

Dossey, A.T., Whitaker, J.M., Dancel, M.C.A., Vander Meer, R.K., Bernier, U.R., Gottardo, M. and Roush, W.R., 2012. Defensive spiroketals from Asceles glaber (Phasmatodea): absolute configuration and effects on ants and mosquitoes. Journal of chemical ecology, 38(9), pp.1105–1115.

Gill, S.A. and Haggerty, T.M., 2012. A comparison of life-history and parental care in temperate and tropical wrens. Journal of Avian Biology, 43(5), pp.461–471.

Guilford, T., 1990. Evolutionary pathways to aposematism. Acta Oecologica-International Journal of Ecology, 11(6) 835–841.

Hansen, B.T., Holen, Ø.H. and Mappes, J., 2010. Predators use environmental cues to discriminate between prey. Behavioral Ecology and Sociobiology, 64(12), pp.1991–1997.

Healy, P.J., 1969. Studies on poisoning by macrozamia communis—I: Biochemical disturbances in the liver. Biochemical Pharmacology, 18(1), pp.85–92.

Henne, P., Kulesza, A., Perez, K. and Houcek, A., 2021. Counterfactual thinking and recency effects in causal judgment. Cognition, 212, p.104708.

Holen, Ø.H. and Svennungsen, T.O., 2012. Aposematism and the handicap principle. The American Naturalist, 180(5), pp.629–641.

Kikuchi, D.W., Allen, W.L., Arbuckle, K., Aubier, T.G., Briolat, E.S., Burdfield-Steel, E.R., Cheney, K.L., Daňková, K., Elias, M., Hämäläinen, L. and Herberstein, M.E., 2023. The evolution and ecology of multiple antipredator defences. Journal of Evolutionary Biology, 36(7), pp.975–991.

Jones, R., C. Davis, S. and Speed, M.P., 2013. Defence cheats can degrade protection of chemically defended prey. Ethology, 119(1), pp.52–57.

Leimar, O., Enquist, M. and Sillen-Tullberg, B., 1986. Evolutionary stability of aposematic coloration and prey unprofitability: a theoretical analysis. The American Naturalist, 128(4), pp.469–490.

Malcolm, S.B. and Brower, L.P., 1989. Evolutionary and ecological implications of cardenolide sequestration in the monarch butterfly. Experientia, 45(3), pp.284–295.

Mathis, A. and Unger, S., 2012. Learning to avoid dangerous habitat types by aquatic salamanders, *Eurycea tynerensis*. Ethology, 118(1), pp.57–62.

McElreath, R., 2018. Statistical rethinking: A Bayesian course with examples in R and Stan. chapman and hall/cRc. Ohno, S. 1970. Evolution by Gene Duplication. Springer-Verlag, Berlin, Heidelberg, New York.

Paul, V.J. and Van Alstyne, K.L., 1988. Use of ingested algal diterpenoids by Elysia halimedae Macnae (Opisthobranchia: Ascoglossa) as antipredator defenses. Journal of Experimental Marine Biology and Ecology, 119(1), pp.15–29.

Poulton, E.B., 1890. The colours of animals: their meaning and use, especially considered in the case of insects (Vol. 67). D. Appleton, New York City, New York

R Core Team. 2023. R: A Language and Environment for Statistical Computing. R Foundation for Statistical Computing, Vienna, Austria. https://www.R-project.org/.

Rothschild, M., Nash, R.J. and Bell, E.A., 1986. Cycasin in the endangered butterfly *Eumaeus atala florida*. Phytochemistry, 25(8), pp.1853–1854.

Rozin, P., Haidt, J., & McCauley, C. R. 2008. Disgust. In Handbook of emotions (3rd ed., pp. 757–776). The Guilford Press.

Ruxton, G.D., Allen, W.L., Sherratt, T.N. and Speed, M.P., 2019. Avoiding attack: the evolutionary ecology of crypsis, aposematism, and mimicry. Oxford University Press

Santos, J.C., Coloma, L.A. and Cannatella, D.C., 2003. Multiple, recurring origins of aposematism and diet specialization in poison frogs. Proceedings of the National Academy of Sciences, 100(22), pp.12792–12797.

Sarabian, C., Wilkinson, A., Sigaud, M., Kano, F., Tobajas, J., Darmaillacq, A.-S., Kalema-Zikusoka, G., Plotnik, J. M., & MacIntosh, A. J. J. 2023. Disgust in animals and the application of disease avoidance to wildlife management and conservation. Journal of Animal Ecology, 92, 1489–1508.

Sherratt, T.N., 2003. State-dependent risk-taking by predators in systems with defended prey. Oikos, 103(1), pp.93–100.

Sherratt, T.N. and Beatty, C.D., 2003. The evolution of warning signals as reliable indicators of prey defense. The American Naturalist, 162(4), pp.377–389.

Skelhorn, J. and Rowe, C., 2010. Birds learn to use distastefulness as a signal of toxicity. Proceedings of the Royal Society B: Biological Sciences, 277(1688), pp.1729–1734.

Speed, M.P. and Ruxton, G.D., 2005. Aposematism: what should our starting point be?. Proceedings of the Royal Society B: Biological Sciences, 272(1561), pp.431–438.

Stan Development Team. 2025. RStan: the R interface to Stan. R package version 2.37.x. https://mc-stan.org/.

Stanback, M.T. and Burke, T.H., 2020. Neophobia in common feeder birds of a southeastern suburb. Southeastern Naturalist, 19(2), pp.333–338.

Stephens, D.W. and Krebs, J.R., 1986. Foraging theory (Vol. 6). Princeton university press.

Strzelewicz, M.A., Ullrey, D.E., Schafer, S.F. and Bacon, J.P., 1985. Feeding insectivores: increasing the calcium content of wax moth (*Galleria mellonella*) larvae. The Journal of Zoo Animal Medicine, 16(1), pp.25–27.

Turner, J.R., Kearney, E.P. and Exton, L.S., 1984. Mimicry and the Monte Carlo predator: the palatability spectrum, and the origins of mimicry. Biological Journal of the Linnean Society, 23(2-3), pp.247-268.

Wasserlauf, Y., Gancz, A., Ben Dov, A., Efrat, R., Sapir, N., Dor, R. and Spiegel, O., 2023. A telemetry study shows that an endangered nocturnal avian species roosts in extremely dry habitats to avoid predation. Scientific Reports, 13(1), p.11888.

